# Physiological stress responses to non-mimetic model brood parasite eggs: leukocyte profiles and heat-shock protein Hsp70 levels

**DOI:** 10.1101/2022.01.31.478544

**Authors:** Francisco Ruiz-Raya, Teresa Abaurrea, Ramón Vigo, Manuel Soler

## Abstract

Obligate avian brood parasites lay their eggs in the nest of other bird species (hosts). Brood parasitism often imposes severe fitness costs on hosts, which selects for the evolution of effective anti-parasitic defences, such as recognition and rejection of brood parasite eggs. Glucocorticoids have been recently found to mediate host physiological and behavioural adjustments in response to brood parasite eggs; however, it remains unclear whether brood parasitism triggers a general response involving multiple physiological elements. In this study, we experimentally investigated whether a salient brood parasitic stimulus (the presence of a non-mimetic model egg in the nest) causes physiological adjustments in adult Eurasian blackbirds (*Turdus merula*) at immune (leukocyte profiles) and cellular (heat-shock protein Hsp70 synthesis) level. Also, we explored whether these physiological changes are mediated by variations in corticosterone levels. We found that experimental brood parasitism caused an increase in heterophils and a decrease in lymphocytes, leading to higher H/L ratios in parasitized birds. Nevertheless, we did not find trade-offs between immune function and corticosterone levels. Hsp70 synthesis was not affected by our experimental manipulation. Our findings provide evidence that brood parasite eggs trigger a general stress response in egg-rejecter hosts, including changes in cellular immune profiles.

## INTRODUCTION

Obligate avian brood parasites, which account for approximately 1% of bird species (Mann, 2017), lay their eggs in the nests of heterospecific birds (hosts), taking advantage of the parental care that hosts provide to their young (Payne, 1977). Interspecific brood parasitism imposes significant fitness costs on hosts, which selects for the evolution of anti-parasitic host defences (e.g., the recognition and rejection of parasitic eggs; Feeney et al., 2014; Soler, 2014) and ultimately can lead to co-evolutionary arms races between brood parasites and hosts (Rothstein, 1990). Over the past decades, much research has focused on the ecological and behavioural aspects of avian brood parasite-host interactions (Soler, 2017). However, the physiological mechanisms underlying host responses to brood parasitism have received comparatively little attention despite the fact that brood parasitism may potentially trigger significate adjustments in host physiology, which can have important consequences for the expression and evolution of key anti-parasitic defences such as egg rejection (Abolins-Abols and Hauber, 2018; Avilés, 2018; Ruiz-Raya, 2021).

Previous studies on host physiological responses to brood parasitism have focused primarily on the endocrinology of stress. Glucocorticoid hormones are known to mediate allostasis in vertebrates, triggering physiological and behavioural changes that help individuals to cope with environmental challenges (Breuner et al., 2008; Wingfield et al., 1998), including avian brood parasitism (Abolins-Abols and Hauber, 2020). Brood parasitism stimuli is known to increase corticosterone (CORT) levels in adult hosts during incubation (Ruiz-Raya *et al*. 2018; but see Scharf *et al*. 2021) and nestling stages (Antonson et al., 2020). Parasitized birds also show elevated CORT responsiveness to stressors during the fledgling period, which can lead to detrimental long-term consequences (Mark and Rubenstein, 2013). Theory predicts that physiological responses to stress will operate at different levels and include multiple physiological systems working together (Wingfield and Romero, 2015). Thus, characterizing the stress response to avian brood parasitism will require assessing different physiological biomarkers that provide supplementary information on the nature of these physiological adjustments (Breuner et al., 2013; MacDougall-Shackleton et al., 2019; O’dell et al., 2014).

A crucial aspect of host physiology that could be affected by brood parasitism is immune function. Environmental stressors are known to cause changes in the relative proportion of white blood cell types (i.e. leukocytes; Davis et al., 2008), a highly conserved physiological response in vertebrates. This parameter has become a widely applied tool in ecophysiology to assess individual responses to stress (Davis et al., 2008; O’dell et al., 2014). The relative proportion of heterophils and lymphocytes (H/L ratio), the two most abundant white cell types in birds, is known to increase in response to external stressors such as climatic conditions, parasites or social challenges (Davis et al., 2008; Minias, 2019; Minias et al., 2018). These stress-induced changes in leukocyte number are typically slower and last longer (from one hour to days) than rapid CORT responses, making leukocyte biomarkers particularly informative for obtaining measures of chronic environmental stress (Davis and Maney, 2018; O’dell et al., 2014). Importantly, short-term changes in H/L ratios may be mediated by glucocorticoids (Sapolsky et al., 2000), although stress hormones and leukocyte profiles are not always correlated (Davis and Maney, 2018). Previous studies have shown that rearing brood parasitic nestlings may cause reduced humoral immune responses in hosts (Antonson et al., 2020), yet the effects of brood parasitism on the components of cell-mediated immunity are still unknown.

Other biomarkers, such as heat-shock proteins (Hsp), in particular the Hsp60 and Hsp70 families, have been widely used to assess long-term chronic stress in wild bird populations (Herring and Gawlik, 2007; O’dell et al., 2014). Hsp are molecular chaperones involved in cellular ‘house-keeping’ functions, whose expression is induced to protect cells from damage caused by a wide range of stressors associated with parasites, environmental or social challenges (Martínez-Padilla et al., 2004; O’dell et al., 2014; Sørensen et al., 2003). This provides valuable supplementary information to hormonal and immune indicators (O’dell et al., 2014). Hsp expression is thought to be part of a general stress response (Asea and Kaur, 2018), and may be associated with glucocorticoid levels (Asea and Kaur, 2018; Mahmoud et al., 2004). The combined use of different biomarkers may therefore help to elucidate the nature and timing of host stress responses; however, there is still little information on the effect of avian brood parasitism on leukocyte profiles and stress protein expression in adult hosts.

Here, we investigate whether a salient brood parasitism stimulus (the presence of one parasitic egg in the nest) triggers significant adjustments in host physiology. Specifically, we evaluated different biomarkers of physiological stress at the immune (leukocyte profile) and cellular level (Hsp expression) in experimentally parasitized and non-parasitized adult hosts. We predict that if the presence of a non-mimetic brood parasite egg induces a general stress response in adult hosts, then we will find elevated H/L ratios and increased Hsp70 expression caused by experimental parasitism. Additionally, we take advantage of our own data on the glucocorticoid response to experimental brood parasitism (from the same individuals, Ruiz-Raya et al., 2018) to explore, through structural equation modelling, whether the effects of experimental brood parasitism on H/L ratios and Hsp70 expression are mediated indirectly by variations in plasma CORT.

## MATERIAL AND METHODS

### Study system

Our study was conducted in a Eurasian blackbird (*Turdus merula*) population located in the Valley of Lecrín, Spain, from March to May 2015. The Eurasian blackbird (*Turdus merula*, hereafter blackbird) is an occasional common cuckoo (*Cuculus canorus*) host frequently used in brood parasitism studies (see e.g., Grim et al., 2011; Roncalli et al., 2019; Ruiz-Raya et al., 2015; Samas et al., 2011; Soler et al., 2015; Soler et al., 2017). Female blackbirds, the sex responsible for egg rejection in this species (Ruiz-Raya et al., 2019), show fine-tuned egg-recognition abilities (see references above).

### Field procedure

From the beginning of the breeding season, we located active blackbird nests, which were visited every two days to obtain data on laying date and clutch size. The day after clutch completion, breeding pairs were randomly selected to incubate clutches either with (parasitized group, n =18) or without non-mimetic parasitic model eggs (non-parasitized control group, n = 16). Following a previously established methodology, parasitic models eggs were painted red to simulate non-mimetic eggs (Avilés et al., 2004; Martín-Vivaldi et al., 2012; Roncalli et al., 2017; Soler and Møller, 1990), which are easily detected by blackbirds (Ruiz-Raya et al., 2019, 2015; Soler et al., 2015). As model eggs, we used natural (commercial) common quail (*Coturnix coturnix*) eggs (32.6 ± 0.1 × 25.3 ± 0.1 mm; n = 49) slightly larger than blackbird eggs (30.4 ± 0.2 × 21.1 ± 0.1 mm; n = 40), a type of model egg previously used to elicit egg recognition in blackbirds (Ruiz-Raya et al., 2018; Soler et al., 2017). In our study population, blackbird clutch size varies from 2 to 5 eggs (Ibáñez-Álamo and Soler 2010), but we only used nests containing 2 or 3 eggs to avoid exceeding the maximum natural clutch size after experimental parasitism. No blackbird ejected the parasitic model egg or deserted the nests by the end of brood parasitism trials.

72 hours after the introduction of the parasitic model egg, all focal females were captured (6:00 – 8:00 am) by using a mist net placed near the focal nest (1 - 5 m). Such 72-hours period has been proved to be a time frame suitable to assess sustained physiological changes in response to experimental brood parasitism (Ruiz-Raya et al., 2018). Immediately after capture (< 3 min), a blood sample (400-500 μl) was collected from the brachial vein with a 25-gauge needle and 80 μl heparinized microhematocrit tubes. Additionally, a drop of blood was transferred to a slide to make one-cell-layer blood smears from both parasitized and control females. Smears were air-dried and stored in darkness until methanol fixation. All females were marked with individual rings and released near the nest 5-15 minutes after blood sampling. In all cases, experimental females returned to the focal nest to resume incubation within the next hour (as revealed by warm clutches). Blood samples were kept cold and, once in the lab, centrifuged at 4500 RCF for 3 min (max. 4 hours after collection). Plasma and red blood cells (RBC) were separated and stored at −20 °C until laboratory assays. Blood smears were fixed in methanol (Houwen, 2002; O’dell et al., 2014).

### Laboratory analyses

Blood smears were stained by using the Giemsa method and scanned, blind to the treatment, at 1000× magnification under a light microscope. Following a general protocol for leukocyte characterization is slides (O’dell et al., 2014), we counted a random sample of 100 leukocytes from each blood smear, and classified them into heterophils (H), lymphocytes (L), and other leukocyte types (i.e., basophils, eosinophils and monocytes) according to the criteria of Hawkey et al., (1989). Then, the H/L ratio was then calculated for each individual by dividing the number of heterophils by the number of lymphocytes. All blood smears were assessed by the same researcher (RV) to reduce variability. Additionally, twenty-five randomly chosen smears were assessed twice to estimate repeatability of H/L ratio measurements, confirming that leukocyte count was highly repeatable (intra-class correlation coefficient, ICC = 0.86, *p* < 0.001). Hsp70 expression was quantified from red blood cells at the Ecophysiology Laboratory of the Estación Biológica de Doñana (Spanish National Research Council, Spain) using a commercial ELISA kit (ADI-EKS-700B, ENZO Biochem Inc., Farmengdale, New York) by following the manufacturer instructions. Total proteins were measured using the Bradford method (Kruger, 1994) and Hsp70 values were corrected according to total protein concentration in the samples. CORT levels were measured from plasma samples by heterologous radioimmunoassay (RIA) following a protocol previously validated for blackbirds (see Ruiz-Raya *et al*. 2018 for additional details on CORT assays).

### Statistical analyses

All analyses and graphs were performed using R version 3.6.1 (R Core Team, 2019). We used linear models (LMs) to assess between-groups differences in heterophil (Box-Cox transformed), lymphocyte, H/L ratio (Box-Cox transformed) and Hsp70 levels. All models included the brood parasitism treatment, the clutch size (two/three) and the two-way interaction between these terms.

Structural equation modeling (SEM) was used to examine direct and indirect causal relationships between our brood parasitism treatment, the main biomarker of the leukocyte response to stress (the H/L ratio; O’dell et al., 2014), and Hsp70 expression by using the *piecewiseSEM* package (Lefcheck, 2016). First, we explored direct links between experimental brood parasitism and heterophils, lymphocytes and Hsp70 expression, as well as indirect paths through the links with plasma corticosterone concentration (full model). The final model was selected by using Shipley’s extension for the Akaike Information Criteria (AIC; Shipley, 2013) and evaluated its goodness of fit using the Fisher’s *C* statistic (Lefcheck, 2016). All models described above satisfied the linearity and homoscedasticity criteria.

## RESULTS

Experimentally parasitized females showed a higher number of heterophiles (F_1,30_ = 8.25, *p* = 0.007, Fig. 1a), and a lower number of lymphocytes (F_1,30_ = 10.60, *p* = 0.004. Fig. 1b), compared to non-parasitized control females. As expected, parasitized females showed a higher H/L ratio than non-parasitized control females (F_1,30_ = 9.11, *p* = 0.005, Figure 1c). Neither the clutch size nor its interaction with the experimental treatment had an effect on the components of the cellular immunity (i.e., heterophil and lymphocyte counts) or the H/L ratio (*p* > 0.27 in all cases). Contrary to our prediction, Hsp70 expression was not affected by our brood parasitism manipulation (F_1,30_ = 0.01, *p* = 0.84, Fig. 1d), independently of clutch size (F_1,30_ = 0.25, *p* = 0.62).

**Figure 1.**
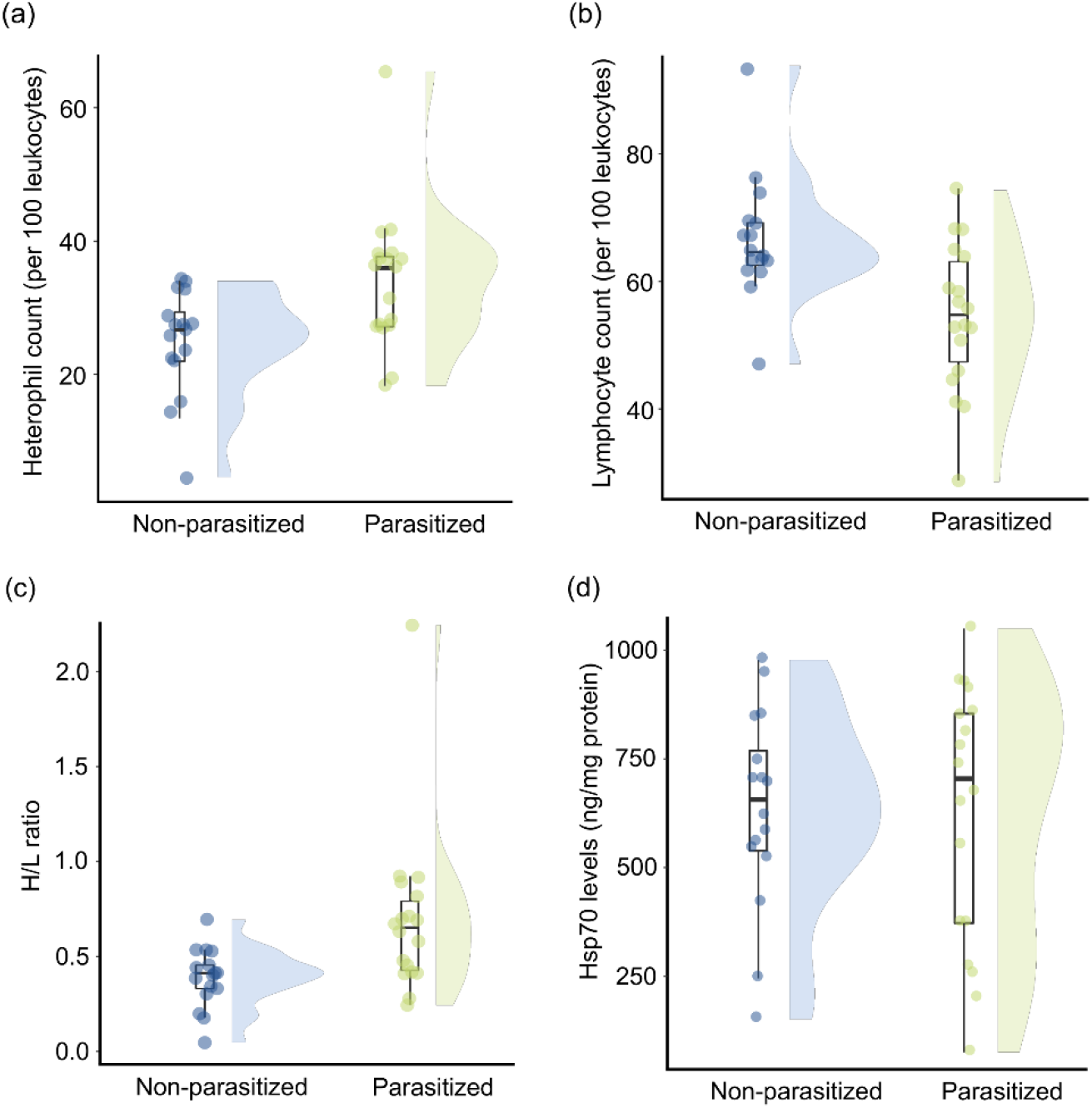
Differences in **(a)** heterophils, **(b)** lymphocites, **(c)** heterophils/lymphocites ratio (H/L ratio) and **(d)** heat-shock protein Hsp70 levels between parasitized and non-parastized control females. Boxplots show the median (bold line), and 25th and 75th percentiles (coloured boxes), with whiskers denoting the 5th and 95th percentiles. The violin plot outlines illustrate the probability density of data, i.e. the width of the shaded area indicates the proportion of the data located there.

SEM analyses confirmed that experimental brood parasitism had a large direct positive effect on the H/L ratio (Fig. 2, Table S1), but no indirect effects via CORT were detected (Fig.2, Table S1). As expected, we found a direct positive effect of the brood parasite stimulus (i.e., the presence of a non-mimetic egg in the nest) on plasma CORT concentration (Fig. 2, Table S1).

**Figure 2.**
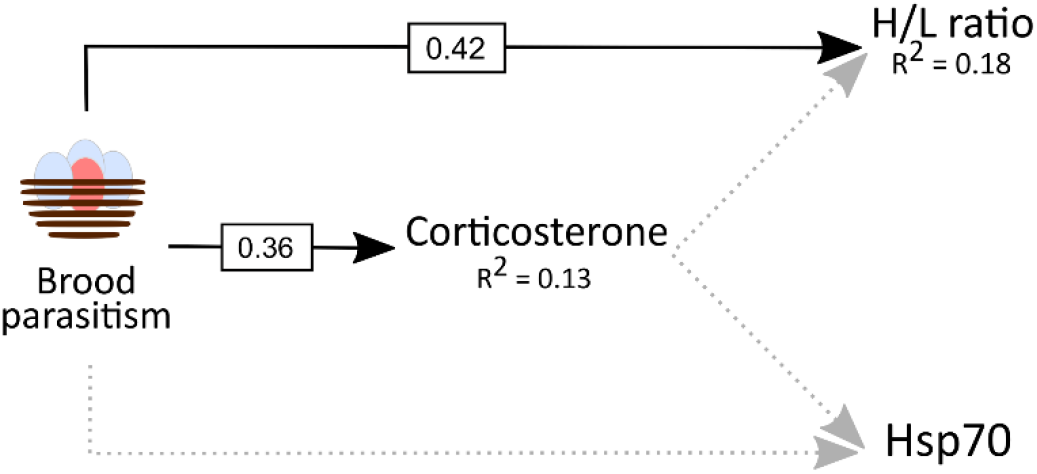
Structural equation model assessing for direct and indirect paths between experimental brood parasitism and the H/L ratio and Hsp70 levels. Plasma corticosterone concentration was included as an indirect path. Grey dotted arrows represent those paths that were tested in the full model but not included in the final model, of which paths are indicated by black arrows. Standardized effects are provided for those paths included in the final model.

## DISCUSSION

We provide evidence that brood parasitic egg stimulus caused significant changes in host leukocyte profiles and, as a result, experimentally parasitized birds showed a higher H/L ratio compared to non-parasitized control individuals. Importantly, these effects were not mediated by plasma glucocorticoid concentration. At the cellular level, our experimental manipulation did not affect the expression of stress proteins. To our knowledge, this is the first evidence of the effects of brood parasitism on the immune status of adult hosts during the incubation phase.

We found that the presence of a non-mimetic model egg in the nest caused a significant increase in heterophils and a decrease in lymphocytes, resulting in higher H/L ratios in parasitized birds (Fig. 1). These physiological adjustments could be caused by different factors related to the presence of parasitic eggs in the nest. First, given the high recognition abilities shown by female blackbirds (see e.g., Samas et al., 2011; Ruiz-Raya et al., 2019), changes in immune function could be part of a general stress response triggered by the recognition of foreign eggs. Changes in host physiology in response to brood parasitism may also include variations in glucocorticoid levels (Ruiz-Raya et al., 2018), which can promote anti-parasitic responses (Abolins-Abols and Hauber, 2020). Indeed, it has recently been shown that experimental brood parasitism either with mimetic or non-mimetic eggs does not lead to changes in the physiology of the prothonotary warbler (*Protonotaria citrea*), an egg-accepter host of the brown-headed cowbird (*Molothrus ater*) (Scharf et al., 2021). This reinforces the idea that these physiological adjustments are, at least partially, triggered by egg recognition.

However, it is also possible that changes in the H/L ratio are related to increased incubation demands associated with increased clutch size (Davis and Maney, 2018; Hanssen et al., 2005). In our study, the effects of experimental brood parasitism on immune biomarkers were not dependent on clutch size, and previous studies have reported that other indicators of physiological stress, such as CORT levels, remain unaffected in hosts naturally parasitized with mimetic eggs (Mark and Rubenstein, 2013). The results described above suggest that incubation demands associated with an additional (parasitic) egg would cause negligible physiological changes in adult hosts *per se*. On the other hand, physiological adjustments triggered by egg recognition and brood enlargement would be expected to act simultaneously during natural brood parasitism events, although some brood parasites may occasionally remove host eggs when visiting target nests (Reboreda et al., 2017). Our study design was unable to assess the separate effects of these factors, so future experimental designs will need to consider alternative manipulations to elucidate the relative importance of egg recognition and brood enlargement in triggering physiological stress responses to brood parasitism, especially in egg-rejecter species with finely tuned egg-recognition abilities.

Regarding the link between immune function and glucocorticoids, our findings confirmed previously published data on the direct positive effects of non-mimetic eggs on plasma CORT of adult hosts (Ruiz-Raya et al., 2018). However, variation in plasma CORT did not mediate an indirect effect of brood parasitism on leukocyte profiles (Fig. 2). This is consistent with previous studies showing that glucocorticoid levels (CORT or cortisol) and leukocyte profiles (H/L ratio) are not always correlated in wild vertebrates (reviewed in Davis and Maney, 2018). Individual trade-offs between CORT and immune responses (humoral immunity) also appear to be absent in cowbird hosts rearing parasite chicks (Antonson et al., 2020). The lack of correlation between these two measures of physiological stress may be due to differences in the timing of CORT and leukocyte responses to chronic stressors (Davis and Maney, 2018). Thus, it may be plausible that, while leucocyte responses to brood parasite model eggs may persist for relatively long periods, CORT response decline over time in some individuals.

Finally, experimental brood parasitism did not elicit differential physiological responses in terms of Hsp70 levels within three days, and Hsp concentration was not related to variation in CORT levels. The short-term stress associated with brood parasite model eggs during this period of time does not appear to cause rapid up-regulation of stress proteins, whose synthesis is a reliable indicator of chronic stress (O’dell et al., 2014). Nevertheless, we cannot rule out that the expression of Hsp proteins may be affected in scenarios where brood parasitism is expected to involve sustained stress for adult hosts, for example, during rearing of brood parasite nestlings or fledglings.

In conclusion, our results show that the presence of a non-mimetic brood parasite egg in the nest causes significant changes in the cellular immune profiles of adult hosts. These results, together with previous studies on the glucocorticoid response to brood parasite eggs (Ruiz-Raya et al., 2018), as well as evidence from the nestling and fledgling periods (Antonson et al., 2020; Mark and Rubenstein, 2013), indicate that parasitism triggers a generalized stress response affecting multiple physiological components in adult hosts. We encourage the use of different physiological biomarkers in order to gain a comprehensive view of the host physiological response to avian brood parasitism.

## Ethical approval

We performed the study following all relevant Spanish national (Decreto 105/2011, 19 de Abril) and regional guidelines. No female deserted their nest during the 3 days after to our experimental manipulation and none exhibited any long-term effects of the study.

## Acknowledgments

We thank to Jordi Figuerola and Francisco Miranda for their work with Hsp70 analyses, and Olivier Chastel and Charline Parenteau for their work with corticosterone assays. We also thank Gianluca Roncalli and Juan Diego Ibáñez-Álamo for their help during the fieldwork.

## Author contribution

FRR and MS conceived and designed the study. FRR and TA conducted the field work. FRR and RV performed the laboratory work. FRR conducted the data analysis and wrote the first draft. All authors critically contributed to drafts and gave final approval for publication.

