## Supplementary Table 1 for "Physiological stress responses to non-mimetic model brood parasite eggs: leukocyte profiles and heat-shock protein Hsp70 levels"

**Table S1.** Structural equation models (SEMs) testing the effects of the presence of a non-mimetic model egg (path 1: Brood parasitism) and plasma corticosterone levels (path 2: Corticosterone). Significant *p*-values are highlighted in bold.

| | | **Full model** | | **Final model** | |
|---|---|---|---|---|---|
| *Response variable* | *Path tested* | *Estimate ± se* | *p-value* | *Estimate ± se* | *p-value* |
| H/L ratio | Brood parasitism | 0.25 ± 0.13 | **0.052** | 0.31 ± 0.12 | **0.013** |
| | Corticosterone | 0.04 ± 0.03 | 0.194 | | |
| Hsp70 | Brood parasitism | 31.20 ± 97.40 | 0.751 | | |
| | Corticosterone | -21.67 ± 20.97 | 0.309 | | |
| Corticosterone | Brood parasitism | 1.68 ± 0.77 | **0.036** | 1.68 ± 0.77 | **0.035** |
| | | AIC 23.26; Fisher's C = 1.27 | | AIC 21.98; Fisher's C = 5.11 | |
